# Surface-induced formation and redox-dependent staining of outer membrane extensions in *Shewanella oneidensis* MR-1

**DOI:** 10.1101/689703

**Authors:** Grace W. Chong, Sahand Pirbadian, Mohamed Y. El-Naggar

## Abstract

The metal-reducing bacterium *Shewanella oneidensis* MR-1 produces extensions of its outer membrane (OM) and periplasm that contain cytochromes responsible for extracellular electron transfer (EET) to external redox-active surfaces, including minerals and electrodes. While the role of multi-heme cytochromes in transporting electrons across the cell wall is well established, their distribution along *S. oneidensis* OM extensions is also thought to allow lateral electron transport along these filaments. These proposed bacterial nanowires, which can be several times the cell length, would thereby extend EET to more distant electron acceptors. However, it is still unclear why these extensions form, and to what extent they contribute to respiration in living cells. Here, we investigate physical contributors to their formation using *in vivo* fluorescence microscopy. While previous studies focused on the display of *S. oneidensis* outer membrane extensions (OMEs) as a response to oxygen limitation, we find that cell-to-surface contact is sufficient to trigger the production of OMEs, including some that reach >100 µm in length, irrespective of medium composition, agitation, or aeration. To visualize the extent of heme redox centers along OMEs, and help distinguish these structures from other extracellular filaments, we also performed histochemical redox-dependent staining with transmission electron microscopy on wild type and cytochrome-deficient strains. We demonstrate that redox-active components are limited to OMEs and not present on other extracellular appendages, such as pili and flagella. We also observed that the loss of 8 functional periplasmic and outer membrane cytochromes significantly decreased both the frequency and intensity of redox-dependent staining found widespread on OMEs. These results will improve our understanding of the environmental conditions that influence the formation of *S. oneidensis* OMEs, as well as the distribution and functionality of EET components along extracellular appendages.

## 1 Introduction

*Shewanella oneidensis* MR-1 is a Gram-negative, facultative anaerobic heterotrophic bacterium with versatile respiratory capabilities: in its quest for energy, it can utilize an array of soluble and insoluble electron acceptors, from oxygen to extracellular solid surfaces such as minerals and electrodes. This ability to couple intracellular reactions to the respiration of external surfaces, known as extracellular electron transfer (EET), allows microbial catalytic activity to be harnessed on the electrodes of bioelectrochemical technologies ranging from microbial fuel cells to microbial electrosynthesis (Nealson, 2017; Schröder and Harnisch, 2017). As an extensively studied model organism for EET, studies of *S. oneidensis* revealed the critical role of periplasmic and outer membrane multi-heme cytochromes in forming extracellular electron conduits that bridge the cell envelope (Beblawy et al., 2018; Beliaev et al., 2001; Chong et al., 2018; Edwards et al., 2018; Myers and Myers, 2001; Richardson et al., 2012). Specifically, the periplasmic decaheme cytochrome MtrA connects through the MtrB porin to the outer membrane decaheme cytochrome MtrC that, along with another decaheme cytochrome OmcA, function as the terminal reductases of external electron acceptors or soluble electron shuttles (Richardson et al., 2012). In addition to this well-established role in directing electron transfer across the cell envelope, the Mtr/Omc components have been recently shown to facilitate long-distance electron transport across the membranes of multiple cells via a redox conduction mechanism thought to arise from a combination of multistep hopping along cytochrome heme chains and cytochrome-cytochrome interactions (Xu et al., 2018).

*S. oneidensis* also forms extensions of the outer membrane and periplasm that include the Mtr/Omc multi-heme cytochromes responsible for EET (El-Naggar et al., 2010; Pirbadian et al., 2014; Subramanian et al., 2018). These outer membrane extensions (OMEs) are proposed to function as bacterial nanowires that also facilitate long-distance EET through redox conduction. However, in contrast to electrode-spanning cells measured by electrochemical gating (Xu et al., 2018), the cytochrome-dependent conductivity of these proposed bacterial nanowires has only been directly assessed under dry, chemically fixed conditions (El-Naggar et al., 2010; Leung et al., 2013). A full understanding of the role of *S. oneidensis* OMEs will therefore require challenging *in vivo* measurements of their specific impact on extracellular respiration and observations of the membrane protein dynamics that allow inter-cytochrome electron exchange and redox conduction (Zacharoff and El-Naggar, 2017).

Beyond the detailed mechanism of electron transport along these structures, additional questions remain regarding the physical and environmental conditions that trigger their formation. The *S. oneidensis* OMEs can extend to several times the cell length, and have been observed with a range of morphologies from chains of interconnected outer membrane vesicles to membrane tubes (Pirbadian et al., 2014). Since early reports suggested that they form in response to electron acceptor limitation, particularly oxygen limitation (Gorby et al., 2006), subsequent studies involving these OMEs have been performed in oxygen limiting conditions (Barchinger et al., 2016; El-Naggar et al., 2010; Pirbadian et al., 2014; Subramanian et al., 2018). However, while the increased expression and production of multi-heme cytochromes under oxygen limiting and anaerobic conditions is well established (Barchinger et al., 2016; Myers and Myers, 1992; Pirbadian et al., 2014), it is not clear if oxygen limitation is the sole contributor to the membrane extension phenotype in *S. oneidensis*. In fact, a recent gene expression study hinted at independent regulatory mechanisms for extending the membrane and localizing the EET proteins (Barchinger et al., 2016). Furthermore, membrane extensions have been reported in multiple organisms under a variety of growth conditions (Benomar et al., 2015; Dubey and Ben-Yehuda, 2011; Galkina et al., 2011; McCaig et al., 2013; Pande et al., 2015; Shetty et al., 2011; Wanner et al., 2008), including those in the form of vesicle chains (Dubey et al., 2016; Pérez-Cruz et al., 2013; Remis et al., 2014; Subramanian et al., 2018; Wei et al., 2014), as is the case for *S. oneidensis*.

It was previously shown that *S. oneidensis* membrane vesicles, which form the basis of OMEs, are redox-active, and that this activity likely stems from the cytochromes present on the purified vesicles (Gorby et al., 2008). The native-state characterization of cytochromes on the OMEs themselves is so far limited to microscopic observations ranging from immunofluorescence (Pirbadian et al., 2014) to electron cryotomography (Subramanian et al., 2018), rather than mapping the activity of the redox centers. The possible association of redox-active components with other extracellular filaments in *Shewanella*, beyond OMEs, also remains largely unexplored. Recent studies in both bacteria and archaea, however, have demonstrated that a combination of histochemical heme-reactive staining and electron microscopy can be used to visualize redox-dependent activity of cytochromes that enable functions ranging from mineral oxidation to interspecies electron transfer within methanotrophic consortia (Deng et al., 2018; McGlynn et al., 2015).

This study set out to address some of these outstanding questions regarding *S. oneidensis* OMEs. To determine the conditions underlying OME formation, we designed *in vivo* fluorescence microscopy experiments allowing us to examine the specific role of oxygen limitation and other physical conditions which might influence OME production in *S. oneidensis* MR-1. We find that cell-to-surface contact is sufficient to trigger the formation of *S. oneidensis* OMEs under a wide range of conditions. To assess the extent of cytochrome-dependent redox activity in these structures, we implemented heme-dependent staining with transmission electron microscopy to compare OMEs in wild type and cytochrome-deficient strains. In doing so, we also probed 3 types of extracellular filaments (OMEs, flagella, and pili) for these EET components. We find that periplasmic and outer membrane cytochromes are responsible for most of the redox activity detected using this assay, and that these components are limited to OMEs and do not associate with flagella or pili.

## 2 Materials and Methods

### 2.1 Cell Cultivation

For experiments probing the conditions of OME formation with fluorescence microscopy, *S. oneidensis* MR-1 cells were grown aerobically from frozen (−80°C) stock in 50 mL LB broth overnight at 30°C and 150 rpm up to late logarithmic phase (OD_600_ 2.4-2.8). From this overnight culture, 5 mL of cells were collected by centrifugation at 4226 × *g* for 5 min and washed twice in sterile defined medium (Pirbadian et al., 2014). Cells were then introduced into a perfusion flow imaging platform described previously (Pirbadian et al., 2014) or the coverslip-bottom glass reactor described below after appropriate dilution to achieve a desirable cell density on the surface for fluorescence time-lapse imaging.

Heme staining and transmission electron microscopy were performed on anaerobic cultures of *S. oneidensis* MR-1 and JG1486 (ΔMtr/Δ*mtrB*/Δ*mtrE*) (Coursolle and Gralnick, 2012). For both strains, 5 mL of an aerobic overnight LB pre-culture was pelleted by centrifugation, washed in defined medium (Pirbadian et al., 2014), and used to inoculate 100 mL of anoxic defined medium in sealed serum bottles with 30 mM fumarate as the sole electron acceptor. After 24 h at 30°C and 150 rpm, at OD_600_ 0.28, this anaerobic culture was harvested by centrifugation at 7142 × *g* for 10 min, washed by centrifugation (4226 × *g* for 5 min), and resuspended in defined medium for a total volume of 10 mL. Cells were then injected into the perfusion flow imaging platform containing an electron microscopy grid (Subramanian et al., 2018).

### 2.2 Fluorescence Microscopy

In all experiments, the membrane stains FM 4-64FX (Life Technologies; 0.25 µg/mL), FM 1-43FX (Life Technologies; 0.25 µg/mL) or TMA-DPH (Cayman Chemical Company; 10 µM) were used to visualize cells and OMEs on an inverted microscope (Nikon Eclipse Ti-E) using the TRITC, FITC or DAPI channels (Nikon filter sets G-2E/C, B-2E/C, and UV-2E/C) with 500 ms, 500 ms, and 100 ms exposure times, respectively. FM 4-64FX was generally used as the membrane stain, except in experiments with no flow or agitation, as this concentration of dye faded more quickly over time in unmixed solutions.

Two experimental platforms were used for fluorescence imaging experiments: a perfusion flow setup used previously (Pirbadian et al., 2014; Subramanian et al., 2018) or a coverslip-bottom glass reactor constructed to allow gas injection and measurement of dissolved oxygen levels while visualizing cells. The reactor consisted of a clean glass tube (thickness 1.5 mm, interior diameter 24.7 mm, and length 50 mm) glued on to a clean 43 mm × 50 mm no. 1 thickness glass coverslip (Thermo Scientific) using waterproof silicone glue (General Electric). The autoclaved reactor was placed on the inverted microscope, and a peristaltic pump (Cole-Parmer Masterflex L/S Easy-Load II) was used to control injection of filtered air at a rate of 3.6 mL/min into the reactor. The air inlet (22G 3” sterile needle) was placed 1-2 mm from the coverslip bottom of the reactor so as to ensure oxygen availability and good mixing near the focal plane. Time-lapse imaging was started immediately following introduction of 10 mL of the cell-media mixture into the reactor and continued for 2 h with images acquired in 5 min increments. Oxygen levels in the reactor were measured by a dissolved oxygen probe (Milwaukee Instruments MW600) at various levels (e.g. 1 mm from bottom, middle, and 1 mm from top) over time after cells were added. To check whether the planktonic cells also displayed OMEs, imaging was stopped after the surface-attached cells produced OMEs, and 400 µL of the planktonic mixture (obtained within 1-2 mm from the top solution-air interface) was gently pipetted to a new clean coverslip, and immediately imaged for another 2 h.

### 2.3 Heme-Reactive Staining and Transmission Electron Microscopy

All heme staining experiments were performed on cells attached to electron microscopy grids recovered from the perfusion flow imaging platform after confirmation of OME production using fluorescence microscopy (Subramanian et al., 2018). To accomplish this, an X-thick holey carbon-coated, R2/2, 200 mesh Au NH2 London finder Quantifoil EM grid was glued to the glass coverslip, with the carbon film-coated side facing away from the glass, before sealing the perfusion chamber. The chamber was filled with flow medium, then 400-600 µL of washed cells were injected for a surface density of 50-150 cells visible per 74 µm × 74 µm square in the 200 mesh grid. Cells were allowed to settle for 5-15 min on the grid before resuming perfusion flow at a volumetric flow rate of 6.1 ± 0.5 µL/s. Imaging continued for about 3.5 hours in 5-min increments before medium flow was stopped and the chamber opened under sterile medium. The EM grid was then removed, chemically fixed, and prepared for electron microscopy visualization of heme iron, using a staining protocol adapted from (McGlynn et al., 2015). First, the sample was fixed for 30 min in 2.5% glutaraldehyde (dissolved in 25 mM HEPES, pH 7.4, 17.5 g/L NaCl), washed 5 times by soaking 1 min each in buffer (50 mM HEPES, pH 7.4, 35 g/L NaCl), then incubated for 1 h or 2.5 h with the heme-reactive stain 3,3’-diaminobenzidine (DAB; 0.0015 g/mL, dissolved in 50 mM Tris HCl, pH 8) with or without 0.02% hydrogen peroxide (H_2_O_2_). After 5 washes (100 mM HEPES, pH 7.8), the sample was stained for 1 h in 1% osmium tetroxide, and washed again 5 times. The sample was negative stained in 1% uranyl acetate or 1% phosphotungstic acid for 2 min and air dried overnight. Dried samples were stored in a desiccator before transmission electron microscopy (TEM) imaging. TEM images were acquired on a JEOL JEM-2100F instrument operated at 200 kV, a FEI Morgagni 268 instrument operated at 80 kV, or a FEI Talos F200C instrument operated at 200 kV.

To determine and quantify the extent of cytochrome-reactive staining after treatment with DAB, ImageJ was used to measure the mean pixel intensity (arbitrary gray value units reflecting electron transmission) across an area in the interior of an extension (*A*), or an area in the background (*B*). For each image, a background threshold value (*C*) was generated by taking the mean background intensity (*B*) and subtracting its standard deviation (*D*); thus, *C* = *B* – *D*. If the mean intensity of an extension (*A*) was lower than this threshold (*C*), then it was categorized as stained. For each condition (wild type, mutant, and chemical control), the percentage of stained OMEs (*E*) was calculated. To calculate the staining intensity of a single OME (*F*), the mean pixel intensity of the extension (*A*) was subtracted from that of the background (*B*), giving *F* = *B* – *A*. A value of *F* was calculated for each of the OMEs assessed in each replicate experiment for each condition (wild type, mutant, and chemical control). For each condition, the mean of all *F* values was calculated, giving *G*_*WT*_, *G*_*mutant*_, and *G*_*control*_. Then, *G*_*WT*_ and *G*_*mutant*_ were corrected by subtracting *G*_*control*_, where *G*_*WT*_ – *G*_*control*_ = *H*_*WT*_, and *G*_*mutant*_ – *G*_*control*_ = *H*_*mutant*_. These values *H*_*WT*_ and *H*_*mutant*_ represent mean staining intensities of all the OMEs in each strain, corrected for the contribution of negative staining (*G*_*control*_). To calculate the fold difference in staining frequency between wild type and the mutant, the percentage of OMEs stained in the wild type (*E*_*WT*_) was divided by that of the mutant (*E*_*mutant*_). To calculate the fold difference in staining intensity between wild type and the mutant, the mean staining intensity of the wild type (*H*_*WT*_) was divided by that of the mutant (*H*_*mutant*_).

## 3 Results and Discussion

### 3.1 Surface Contact is Sufficient to Induce Production of Outer Membrane Extensions by *Shewanella oneidensis* MR-1

Production of OMEs by a majority of *S. oneidensis* cells was observed in the oxygen limiting perfusion flow platform, as previously described (Pirbadian et al., 2014; Subramanian et al., 2018) (**Fig. 1**), but also in near-saturating oxygen conditions (6.5-7.5 ppm O_2_) provided by a glass-bottomed reactor that allowed air injection during *in vivo* microscopy (**Fig. 2**). Though it can take up to several hours for a majority of surface-attached cells to produce OMEs, we can observe production of OMEs as early as 10 min after cells contact the surface of a glass coverslip (**Figs. 2**, **S1**). To further examine the role of surface contact, planktonic cells from the bulk oxygenated reactor were sampled 2 h after the reactor was inoculated (approx. 1.5 h after OMEs started being produced by surface-attached cells) and transferred to clean coverslips for observation. These previously planktonic cells showed no evidence of OMEs at the time of sampling, but then also went on to begin to display OMEs within 35 min after contacting the surface (**Fig. 2**). These observations were not limited to the defined minimal medium used, a particular surface chemistry, or mixing conditions; post-attachment OME production was also observed in rich (LB) medium or in buffer (PBS), on different surfaces (glass coverslips and carbon-coated electron microscopy grids), and regardless of liquid flow or agitation (**Figs. S1, S2**). To ensure that the used cell density did not result in O_2_-limiting conditions selectively at the surface, we also experimented with sparse coverage, down to 5-20 cells per field of view (112 µm × 112 µm) in a well-mixed and oxygenated reactor, and confirmed that these cells also produced OMEs (**Fig. S2C**).

**Figure 1.**
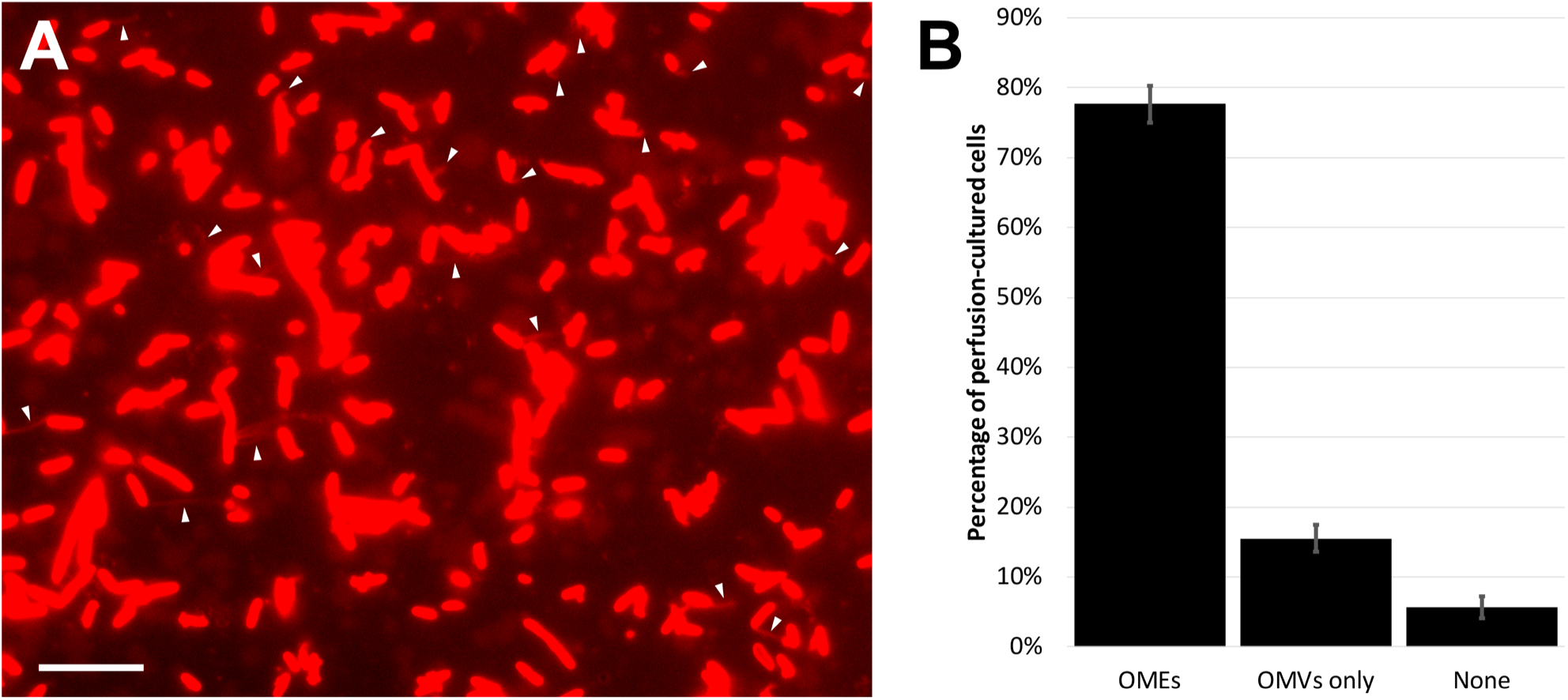
Outer membrane extensions are commonly formed by surface-attached perfusion culture cells. **(A)** Time-lapse fluorescence microscopy snapshot of outer membrane extensions (OMEs, white arrows) produced by *S. oneidensis* MR-1 at a single timepoint in a 3.5-h perfusion flow imaging experiment. Cells and OMEs are visualized with the red membrane stain FM 4-64FX. **(B)** Statistics of OME production from over 5400 cells in 4 replicate 3.5-h perfusion culture experiments illustrates that a majority (78%) of cells produce OMEs visible over time. The remaining cells were seen with only outer membrane vesicles (OMVs), or nothing at all. Error bars show mean ± SEM. (Scale bar: 10 µm.)

**Figure 2.**
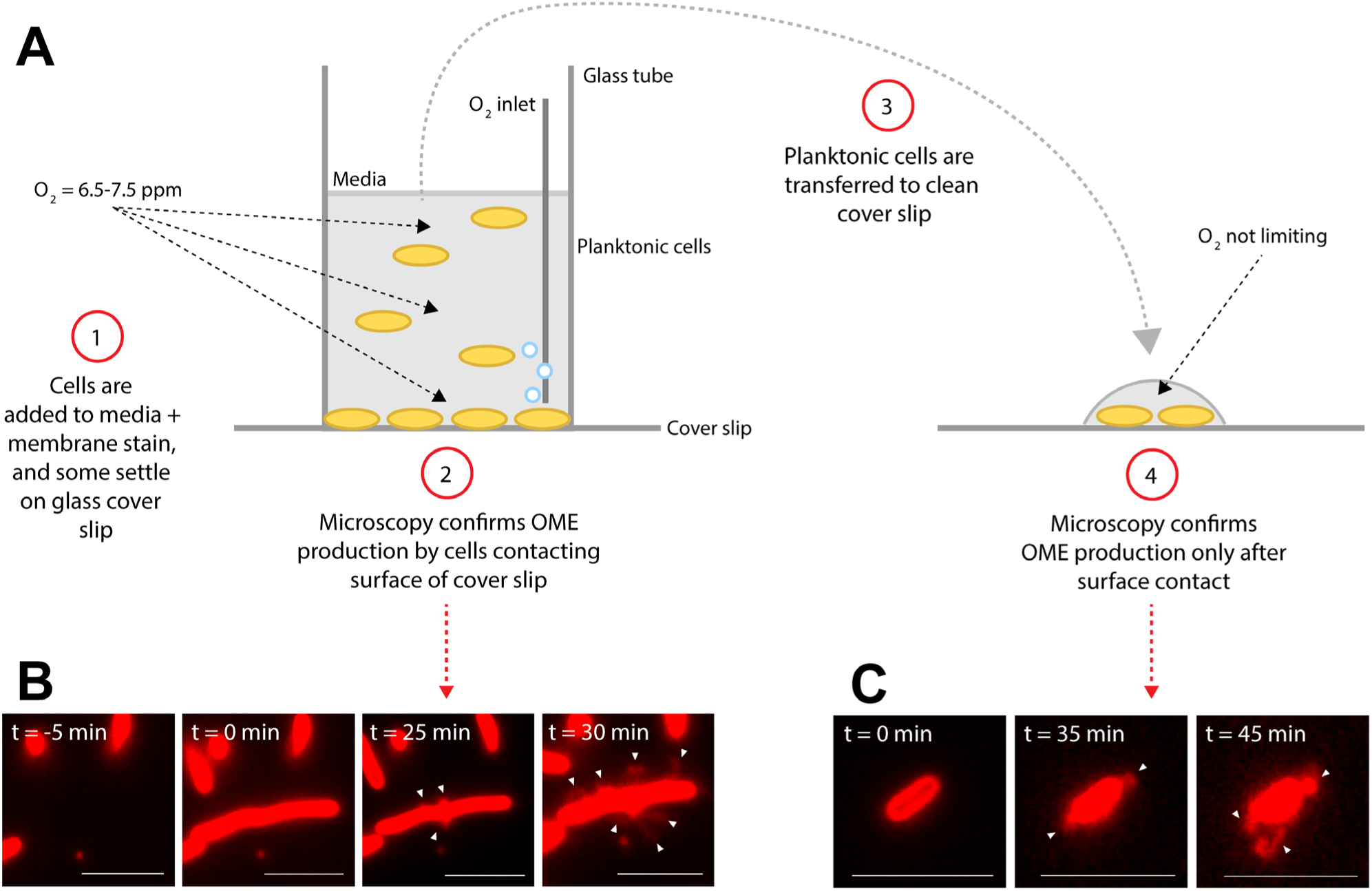
Surface attachment is sufficient to induce production of outer membrane extensions. **(A)** Diagram illustrates experimental procedure. **(B-C)** Microscopy images of *S. oneidensis* MR-1 cells and membrane extensions (white arrows) labeled with the red membrane stain FM 4-64FX. Time (t = 0 min) indicates estimated time of cells contacting the glass surface. **(B)** Demonstrates production of outer membrane extensions (OMEs) by surface-attached cells in the aerated glass-bottomed reactor. **(C)** Demonstrates OME production by planktonic cells from the reactor which were transferred to a new coverslip surface after events in **(B)** were confirmed. (Scale bars: 5 µm.)

Taken collectively, these observations of OME production by surface-attached cells, but not by planktonic cells until subsequent attachment, and regardless of medium composition, surface type, and oxygen availability, point to surface contact as the primary determinant of OME production by *S. oneidensis*. Previous studies on the role of cytochrome-functionalized OMEs as bacterial nanowires primarily focused on the formation of these structures under O_2_-limited conditions (El-Naggar et al., 2010; Gorby et al., 2006; Pirbadian et al., 2014; Subramanian et al., 2018). Our observations suggest that, while O_2_ limitation is necessary for enhanced production of the multi-heme cytochromes required for EET (Barchinger et al., 2016; Myers and Myers, 1992), the membrane extension phenotype is predominantly controlled by surface attachment. Our findings are consistent with a previous proposal based on transcriptome and mutant analyses (Barchinger et al., 2016) that independent pathways are responsible for producing the EET components and extending the outer membrane, while implicating surface contact in controlling the latter pathway.

While our observations show that surface attachment is sufficient to induce OMEs, it is important to note that we do not rule out the influence of O_2_ limitation on the frequency of OME production. In perfusion flow imaging, we are able to precisely define the percentage of OME-producing cells: observation of 5400 cells over four replicate experiments revealed that 78% of surface-attached cells produced OMEs during 3.5 h of perfusion culture (**Fig. 1**). This precise quantification is possible in perfusion flow imaging because the laminar flow helps to restrict the structures to the focal plane near the surface. However, this laminar flow establishes O_2_ limitation as a result of cellular O_2_ consumption and the no-slip condition at the surface-solution interface (Pirbadian et al., 2014). Thus, we could precisely determine the frequency of OME production only in O_2_-limiting perfusion conditions, but not in oxygenated well-mixed reactors where the structures could fluctuate in and out of the focal plane.

Membrane extensions, including those formed as chains of membrane vesicles (MVs), are not limited to *S. oneidensis* (Benomar et al., 2015; Dubey et al., 2016; Dubey and Ben-Yehuda, 2011; Galkina et al., 2011; McCaig et al., 2013; Pande et al., 2015; Pérez-Cruz et al., 2013; Remis et al., 2014; Shetty et al., 2011; Subramanian et al., 2018; Wanner et al., 2008; Wei et al., 2014). The finding that surface contact plays an important role is consistent with prior observations of vesicle chains and OMEs produced by surface-attached cells of other bacteria, including *Shewanella vesiculosa* (Pérez-Cruz et al., 2013), *Bacillus subtilis* (Dubey et al., 2016), and biofilms of *Myxococcus xanthus* (Remis et al., 2014). In addition, another *M. xanthus* study noted that static, rather than shaken, conditions promote more OME production (Wei et al., 2014). The importance of surface-attached, biofilm, or static conditions may point to a generalized mechanism where MVs, which are ubiquitous features of bacteria (Beveridge, 1999; Bohuszewicz et al., 2016; Schwechheimer and Kuehn, 2015), are successively produced and merged into long extensions rather than shed away under more dynamic (e.g. free-swimming or shaken culture) conditions. Once formed, these extensions may then enable a variety of functions ranging from facilitating cell-cell interactions (Dubey et al., 2016; Dubey and Ben-Yehuda, 2011; Remis et al., 2014) to the long-distance EET role proposed for *S. oneidensis* OMEs (El-Naggar et al., 2010; Gorby et al., 2006).

It was also previously proposed that MVs and OMEs can increase the likelihood of encountering neighboring cells and external redox-active surfaces by virtue of the significant change in surface area-to-volume ratio that these structures present (Pirbadian et al., 2014). Consistent with this proposal, we occasionally captured multiple extensions from single cells (**Fig. S3**) as well as *in vivo* fluorescent observations of remarkably long OMEs, likely the longest observed to date. **Fig. 3** and **Movie S1** captures a cell producing a >100 µm OME at a rate over 40 µm/h, at the same time that the cell surface area appeared to shrink by an amount consistent with the newly displayed OME.

**Figure 3.**
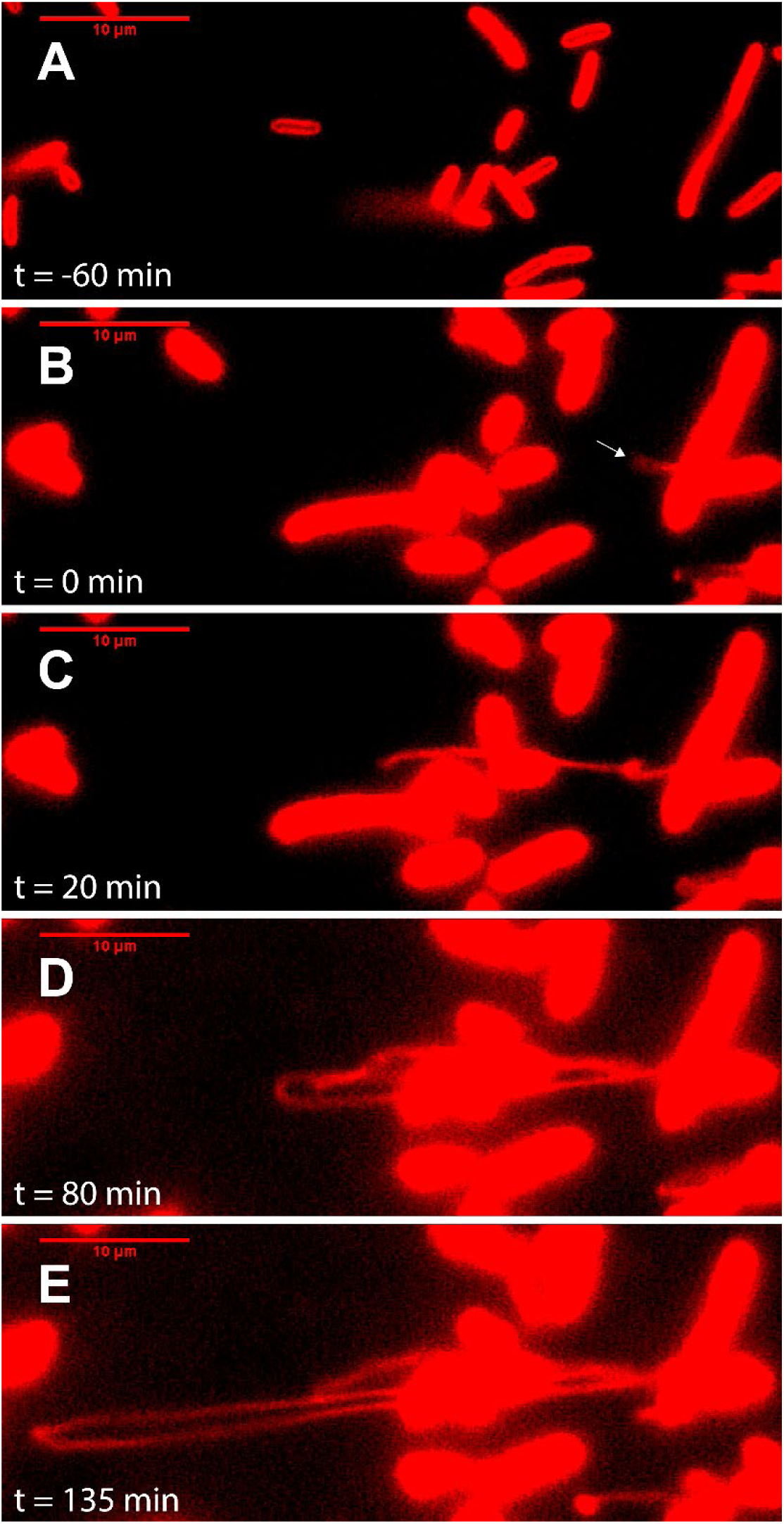
Outer membrane extensions can reach a length of >100 µm, produced at a rate >40 µm/h. Image sequence from time-lapse fluorescence microscopy of surface-attached perfusion cultured *S. oneidensis* MR-1 ΔMtr/Δ*mtrB*/Δ*mtrE* cells. Cells and outer membrane extensions (OMEs) were visualized with the red membrane stain FM 4-64FX. Time (t = 0) marks time since OME production (white arrow). Images depict progression of events, such as **(A)** cell contacting surface, **(B)** OME first visible (white arrow), **(C)** OME elongates and begins to fold, **(D)** further OME elongation, **(E)** OME reaches longest point visible during this experiment. (Scale bars: 10 µm.)

### 3.2 Redox-Dependent Staining of Extracellular Filaments

The localization of the multi-heme cytochromes responsible for EET to OMEs has been previously demonstrated by immunofluorescence observations of MtrC and OmcA (Pirbadian et al., 2014), as well as electron cryotomography observations of outer membrane and periplasmic electron densities consistent with cytochrome dimensions (Subramanian et al., 2018). To examine the distribution and activity of the heme iron redox centers along the OMEs, we applied the heme-reactive 3,3’-diaminobenzidine (DAB)-H_2_O_2_ staining procedure (McGlynn et al., 2015), where the iron centers catalyze the oxidation of DAB, forming a localized dark precipitate that can be observed with the resolution of transmission electron microscopy (TEM). As expected, the OMEs clearly stained for heme, with a noticeable <50 nm band of dark precipitate lining the vesicles that compose the entire structure (**Fig. 4**). Staining was clearly limited to the OMEs and was absent from the other extracellular filaments observed, demonstrating that the cytochromes do not associate with pili and flagella (**Fig. 4**). The absence of staining in these structures, even when observed in contact with the OMEs (**Fig. 4**), also points to the localized nature of the stain. Meanwhile, the <50 nm thickness of precipitate lining OMEs (i.e., precipitate expansion in the direction perpendicular to the surface of the OME) suggests <50 nm lateral distribution of heme redox centers on OMEs, consistent with the surface distribution of putative cytochromes on OMEs visualized by electron cryotomography (Subramanian et al., 2018).

**Figure 4.**
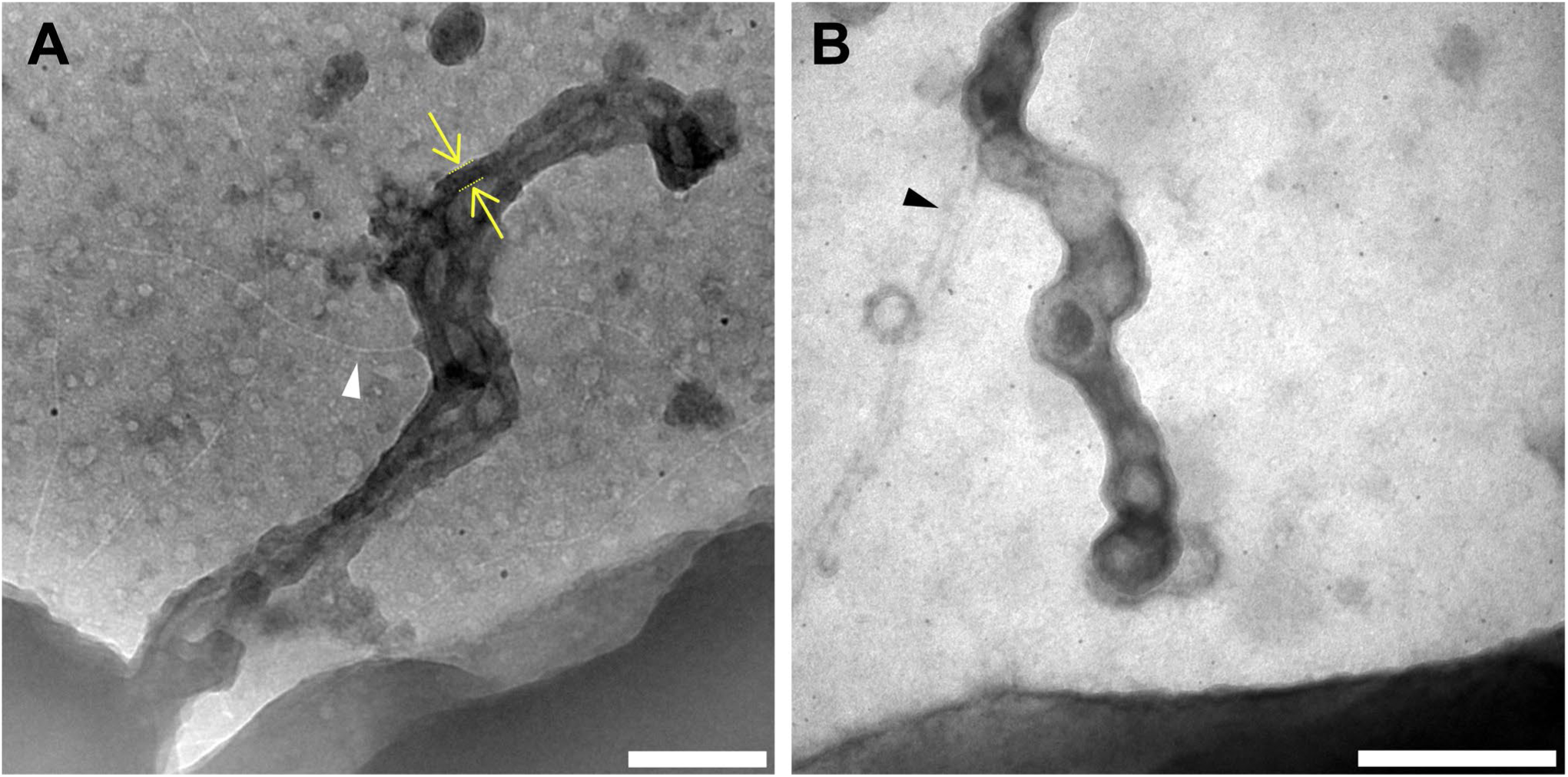
Redox components are present only on outer membrane extensions, not pili or flagella. Histochemical redox-dependent staining with 3,3’-diaminobenzidine (2.5 h staining step) and transmission electron microscopy distinguishes between types of extracellular filaments in *S. oneidensis* MR-1. Images depict dark precipitate (yellow arrows and lines) labeling only outer membrane extensions, but not adjacent extracellular structures **(A)** pili (white arrow) and **(B)** flagella (black arrow). (Scale bars: 200 nm.)

In addition to chemical controls for staining (i.e. wild type with no H_2_O_2_), we also systematically compared OMEs from wild type *S. oneidensis* and a mutant lacking genes encoding eight functional periplasmic and outer membrane cytochromes (ΔMtr/Δ*mtrB*/Δ*mtrE*), including the entire Mtr/Omc pathway of decaheme cytochromes (Coursolle and Gralnick, 2012). This mutant is unable to perform EET (Coursolle and Gralnick, 2012; Wang et al., 2019; Xu et al., 2018) or support long-distance redox conduction across electrodes (Xu et al., 2018). We performed two replicate experiments for each of three conditions (wild type, mutant, and wild type chemical control with no H_2_O_2_), with a total of 45-60 OMEs analyzed per condition. Using image processing to compare OME staining to background intensities (see **Materials and Methods**), we found that the majority (92%) of wild type OMEs stained for heme, but none stained in the chemical control where H_2_O_2_ was omitted (**Fig. 5**). In contrast, a fraction (39%) of OMEs in the mutant strain exhibited heme staining, 2.4-fold less than in wild type (p < 0.0001, Pearson’s chi-squared test) (**Fig. 5**). While lacking all cytochromes necessary for EET, staining in the mutant OMEs is likely due to the additional periplasmic cytochromes, including the flavocytochrome FccA present that functioned as the terminal fumarate reductase to support respiration of fumarate in our anaerobic cultures. Consistent with this interpretation, staining intensity was 3.6-fold stronger in the wild type than in the mutant (p < 0.0001, Student’s *t*-test, two-sample assuming equal variances) (**Fig. 5**). Relative to the mutant control, the observed wild type increase in both staining frequency and intensity indicates that the periplasmic and outer membrane cytochromes necessary for EET contribute much of the redox capacity of the OMEs.

**Figure 5.**
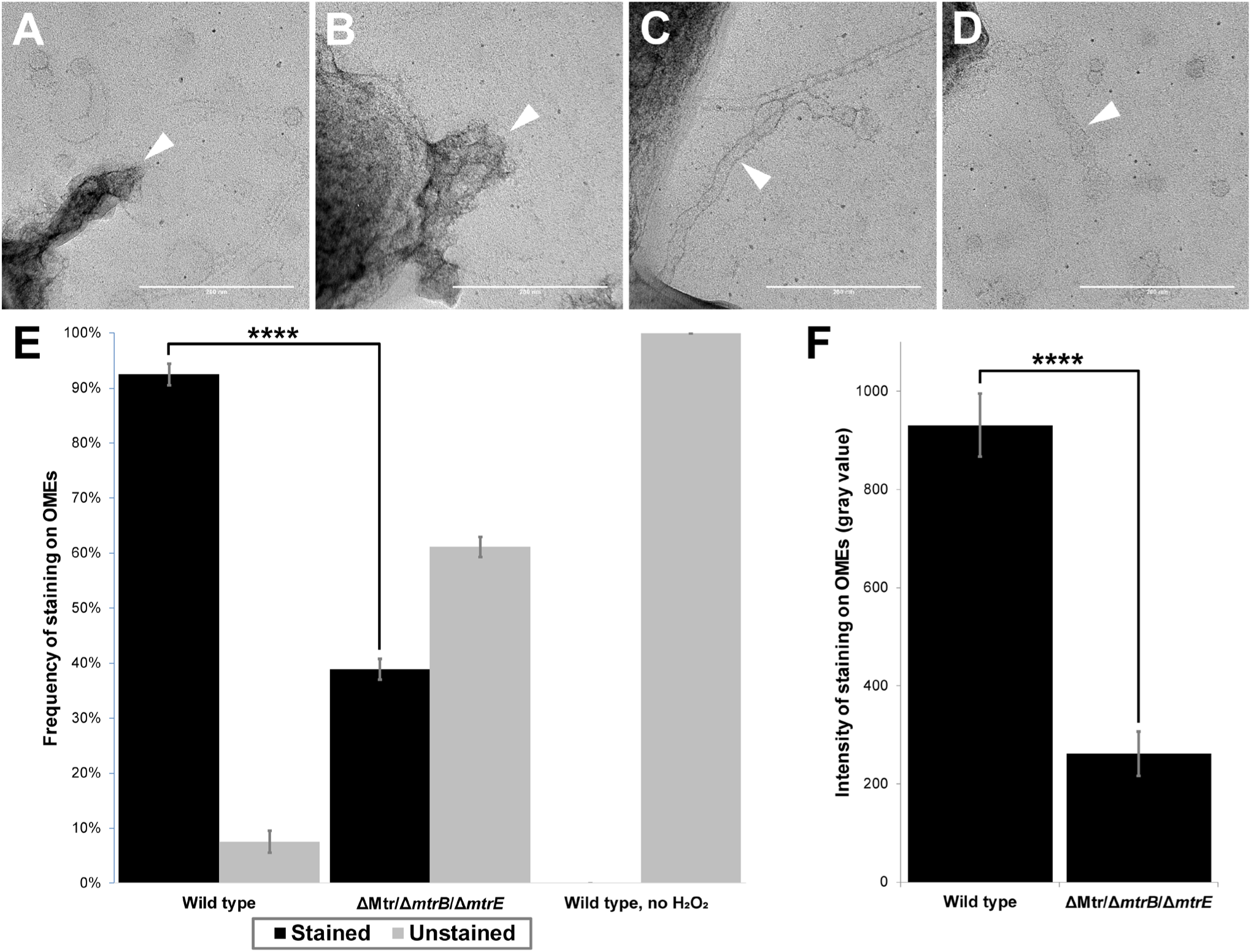
Presence of multi-heme cytochromes important for extracellular electron transfer leads to significantly higher frequency and intensity of redox-dependent staining on outer membrane extensions. **(A-D)** Transmission electron microscopy images depict outer membrane extensions (OMEs, white arrows) stained by 3,3-diaminobenzidine (DAB; 1 h staining step) in wild type and cytochrome-deficient (ΔMtr/Δ*mtrB*/Δ*mtrE*) *S. oneidensis* MR-1 cells. **(A)** Wild type OMEs are stained by DAB precipitate. **(B-C)** Mutant OMEs treated by DAB exhibit varying degrees of staining. **(D)** Wild type OMEs in chemical controls where H_2_O_2_ was omitted appear unstained aside from negative stain. **(E)** Frequency of staining displayed by OMEs in wild type, ΔMtr/Δ*mtrB*/Δ*mtrE* mutant, and wild type chemical control where H_2_O_2_ was omitted. 2.4-fold more OMEs were stained in wild type than in the mutant (p < 0.0001). Statistical significance was determined by p-value from Pearson’s chi-squared test. **(F)** Intensity of staining displayed by OMEs is 3.6-fold higher in wild type than in ΔMtr/Δ*mtrB*/Δ*mtrE* mutant (p < 0.0001). Statistical significance was determined by two-tailed p-value from Student’s *t*-test, two-sample assuming equal variances. Error bars show mean ± SEM. (Scale bars: 200 nm.)

Given its ability to discriminate between cytochrome-containing and cytochrome-free extracellular filaments, and to examine the effect of specific mutations, this heme visualization strategy may hold promise for understanding the presence of redox centers in a variety of microbial systems. However, a detailed understanding of the extent to which these redox centers enable long-distance electron transport along OMEs requires: (i) applying electrochemical techniques, recently used to measure redox conduction in biofilms (Xu et al., 2018; Yates et al., 2016), specifically to OMEs or their MV constituents; and (ii) measurements of the diffusive dynamics of redox molecules along membranes, to test the hypothesis that these dynamics facilitate a collision-exchange mechanism of inter-protein electron transport over micrometer length scales (Subramanian et al., 2018). We are actively pursuing these electrochemical and dynamics measurements.

## 4 Conclusions

In summary, we investigated physical contributors to the production of OMEs by *Shewanella oneidensis* MR-1 and applied heme-reactive staining to examine the extent of the redox centers along the extensions. While previous studies focused on the role of oxygen limitation in triggering the formation of these structures, we demonstrated that surface contact is sufficient to trigger production of OMEs under a variety of medium, agitation, and aeration conditions. In addition, we show that the multi-heme cytochromes necessary for EET contribute much of the redox-dependent staining widespread on OMEs, and that these EET components do not associate with other extracellular filaments. In addition to describing some reproducible microscopic and histochemical techniques to observe redox-functionalized membrane extensions, these observations motivate additional studies to understand the extent to which *Shewanella oneidensis* OMEs can contribute to EET and long-distance redox conduction.

## 6 Author Contributions

G.W.C. designed, performed, and analyzed experiments with guidance from S.P. and M.Y.E-N. G.W.C. and M.Y.E-N wrote and edited the manuscript, with revisions from S.P.

## 8 Acknowledgments

We are grateful to Professor Jeffrey A. Gralnick for providing the cytochrome mutant strain. Transmission electron microscopy at 200 kV was performed at the University of Southern California’s Core Center of Excellence in Nano Imaging. We also thank Christopher Buser for imaging our electron microscopy samples at 80 kV at the Huntington Medical Research Institutes.

