## Supplementary Figures and Legends for "Surface-induced formation and redox-dependent staining of outer membrane extensions in *Shewanella oneidensis* MR-1"

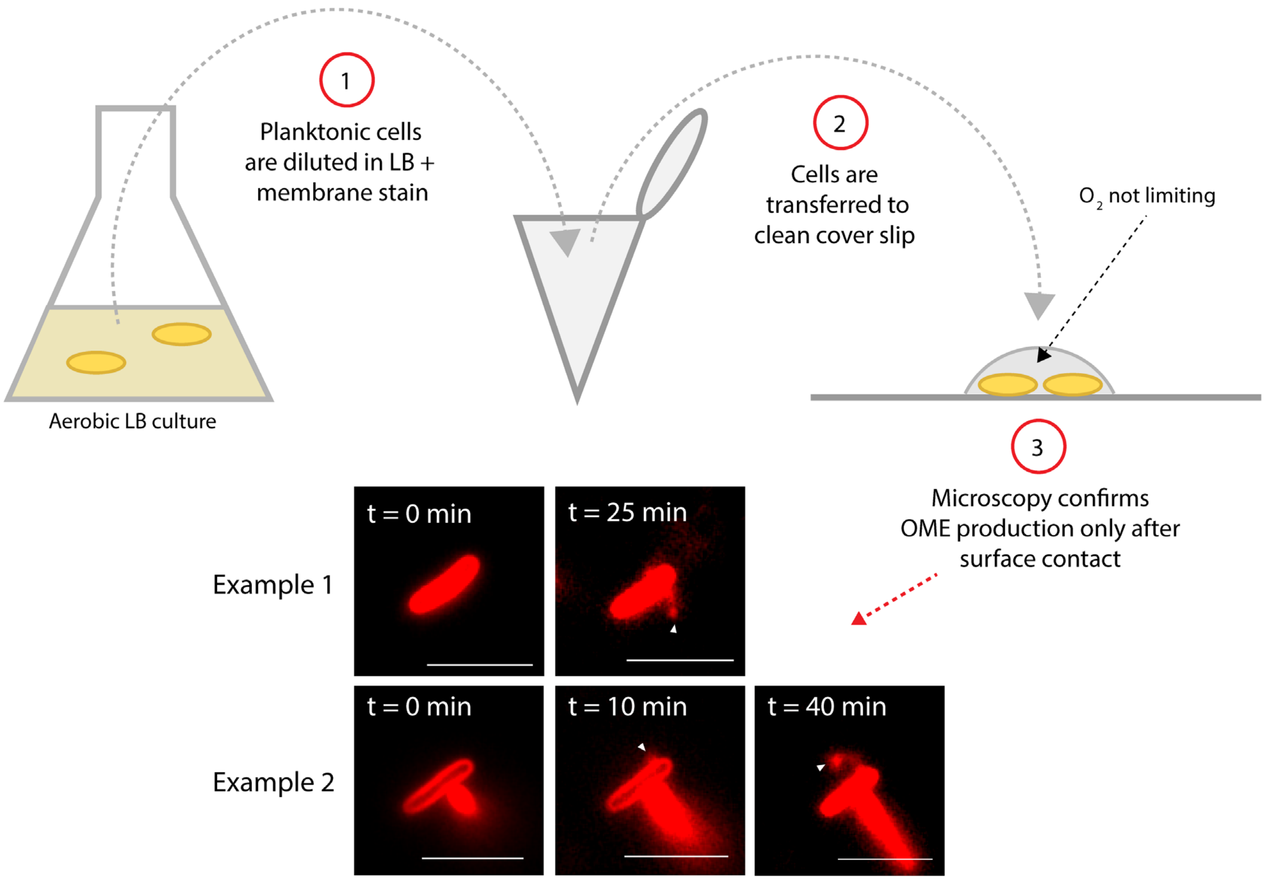


**Figure S1. Outer membrane extensions are produced quickly by planktonic cells in rich aerobic medium soon after cell-to-surface contact.** Diagram illustrates experimental procedure. Microscopy images depict *S. oneidensis* MR-1 cells and outer membrane extensions (OMEs, white arrows) labeled with the red membrane stain FM 4-64FX. Time (t = 0 min) indicates estimated time of cells contacting the glass surface. (Scale bars: 5 µm.)

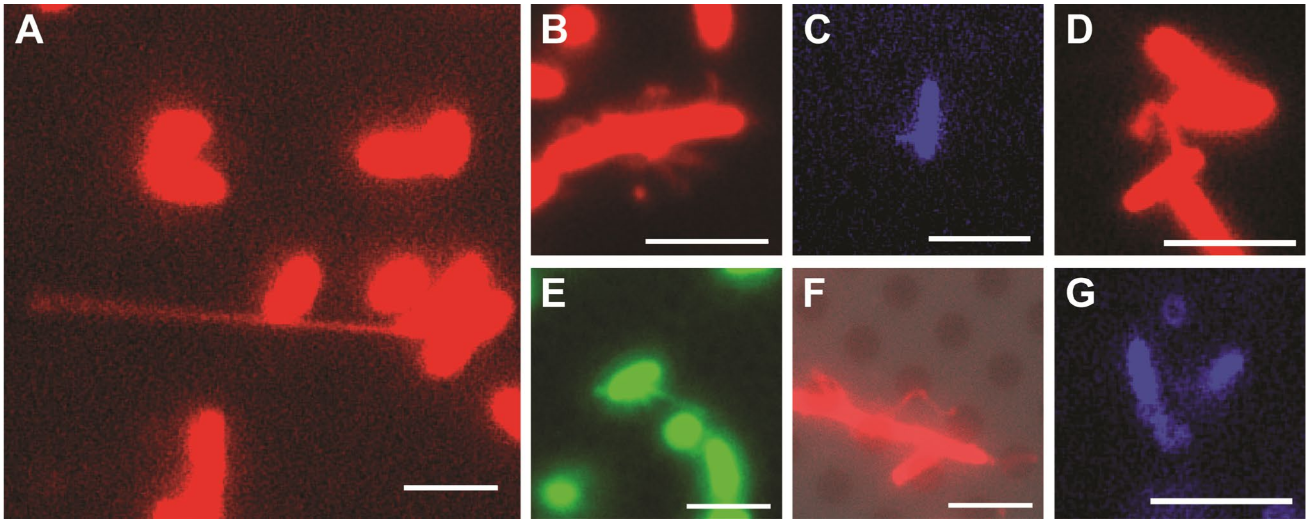


**Figure S2. Outer membrane extensions are produced in a variety of surface-attached conditions, regardless of medium composition, surface chemistry, agitation, or aeration.** *S. oneidensis* MR-1 cells and membrane extensions are visualized by membrane stains FM 4-64FX (red), FM 1-43FX (green), or TMA-DPH (blue). Unless otherwise specified, cells were imaged at the surface of glass coverslips with flow or agitation of oxygen-limited minimal medium. Outer membrane extensions are observed in **(A)** oxygen-limiting perfusion conditions, described previously (Pirbadian et al., 2014; Subramanian et al., 2018), **(B)** oxygen-abundant, high cell density conditions, **(C)** oxygen-abundant, low cell density conditions, **(D)** in rich (LB) medium, **(E)** in buffer (PBS), **(F)** on a carbon-coated electron microscopy grid, and **(G)** without flow or agitation. (Scale bars: 5 µm.)

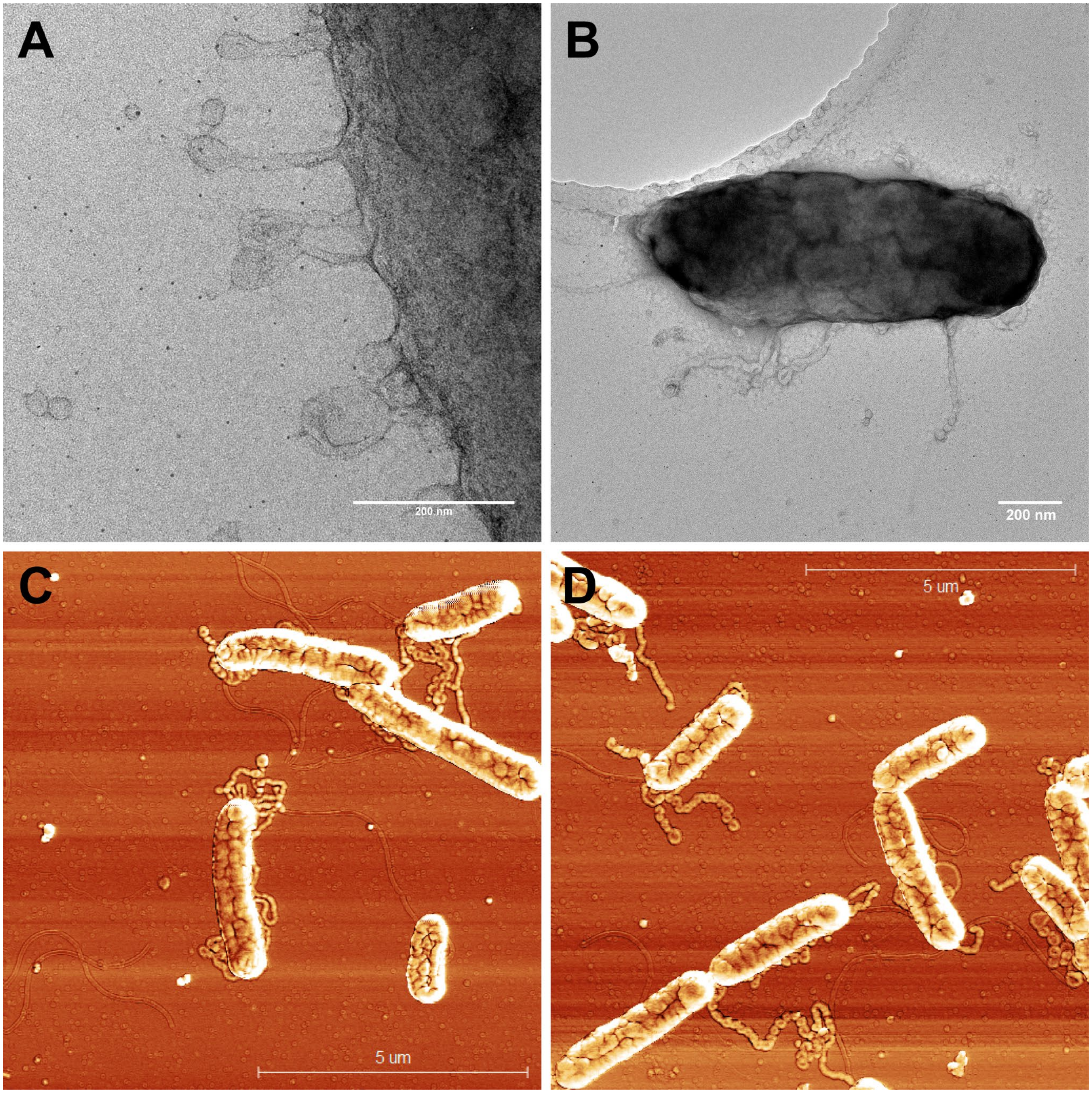


**Figure S3. Cells can produce multiple outer membrane extensions per cell. (A-B)** Transmission electron microscopy images of chemically fixed, negatively stained *S. oneidensis* MR-1 cells and membrane extensions (Scale bars: 200 nm.) **(C-D)** Atomic force microscopy tapping mode phase images of chemically fixed cells and membrane extensions. (Scale bars: 5 µm.)

**Movie S1. Outer membrane extensions can reach a length of >100 µm, produced at a rate >40 µm/h.** Video from time-lapse fluorescence microscopy of surface-attached perfusion cultured *S. oneidensis* MR-1 ΔMtr/Δ*mtrB*/Δ*mtrE* cells. Cells and membrane extensions were visualized with the red membrane stain FM 4-64FX. Images were taken every 5 minutes over 5 hours of perfusion flow. (Scale bar: 10 µm.)
